# Mathematical characterization of fractal river networks

**DOI:** 10.1101/2022.11.07.515375

**Authors:** Akira Terui

## Abstract

Carraro and Altermatt^1^ performed an extensive analysis to study scaling properties of river networks. They concluded that: (1) branching “probability” is better termed as branching “ratio” for real river networks because it is not a probability; (2) branching ratio is a scale-dependent quantity as the value changes across spatial resolutions at which river networks are extracted (expressed as the threshold catchment area *A_T_* that initiates channels or pixel size *l*); (3) Optimal Channel Networks most accurately predicted the metapopulation stability and capacity. However, the supporting ground for these conclusions is either seriously flawed or inconclusive due to the improper use of branching probability, scale invariance, dimensions, and units.

## Maintext

Rivers form complex branching networks, and the ecological implication of river network complexity has gained great interest over the past few decades^2–4^. To this end, there have been concerted efforts to construct virtual river networks to provide theoretical insights into how river network structure controls riverine ecological dynamics. Carraro and Altermatt^1^ compared scaling properties and ecological dynamics in virtual river networks produced by three different simulation methods – balanced binary trees (BBTs)^5^, random branching networks (RBNs)^6,7^, and optimal channel networks (OCNs)^8^. In addition, the authors reported power-law scaling of a structural characteristic (the number of links per unit distance, termed “branching ratio” in their article) in real rivers. Carraro and Altermatt^1^ concluded that: (1) branching “probability” is better termed as branching “ratio” for real river networks because it is not a probability; (2) branching ratio is a scale-dependent quantity as the value changes across spatial resolutions at which river networks are extracted (expressed as the threshold catchment area *A*_*T*_ that initiates channels or pixel size *l*); (3) OCNs most accurately predicted the metapopulation stability and capacity. However, the supporting ground for these conclusions is either seriously flawed or inconclusive due to the improper use of branching probability, scale invariance, dimensions, and units.

First, branching probability *p* is certainly a probability by definition. In previous studies, branching probability *p* has been defined as a cumulative probability distribution of link length *L* [km] in real river networks (a “*link”* represents a river segment from one confluence to another). For example, Terui et al.^6,7^ fitted an exponential distribution to link length as *L_j_* ~ Exp(*θ*) (*θ* is the rate parameter and *j* refers to an individual link) and estimated branching probability as *p* = 1 – exp(–*θL*) (*L* was set to be 1 [km] in Terui et al.^6,7^). Thus, *p* [km^−1^] in this example is the *probability* of including a confluence (or an upstream terminal) per unit river km. Therefore, the author’s claim”…*it is in fact improper to refer to a probability when analyzing the properties of a realized river network*…” is simply a misunderstanding. Instead, branching ratio *p_r_* is identical to the rate parameter *θ* [km^−1^], which is the inverse of average link length 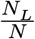 (*N_L_* the number of links in a network [–]; *N* the total river length [km]). To show this equality (*θ* = *p_r_*), I delineated 50 river networks analyzed in Carraro and Altermatt^1^ using MERIT Hydro^9^, with which *θ* and *p_r_* are independently estimated as described above. MERIT Hydro has constant pixel size of 3 arc-second (~90 m at the equator) across the globe. I used *A*_*T*_ = 1 [km^2^] to capture sufficient details of real river networks. The relationship between *p_r_* and *θ* fell exactly on a 1:1 line (Figure 1A), thus *θ* = *p_r_*. By definition, branching probability is a monotonic increasing function of *p_r_* with a possible range of 0 – 1 (Figure 1B); yet, they are not identical. Note that, in the following paragraphs, I use *p_r_* for my analysis to be consistent with the original article. Since branching probability *p* is a simple transformation of *p_r_*, the arguments are translatable between the two.

**Figure 1:**
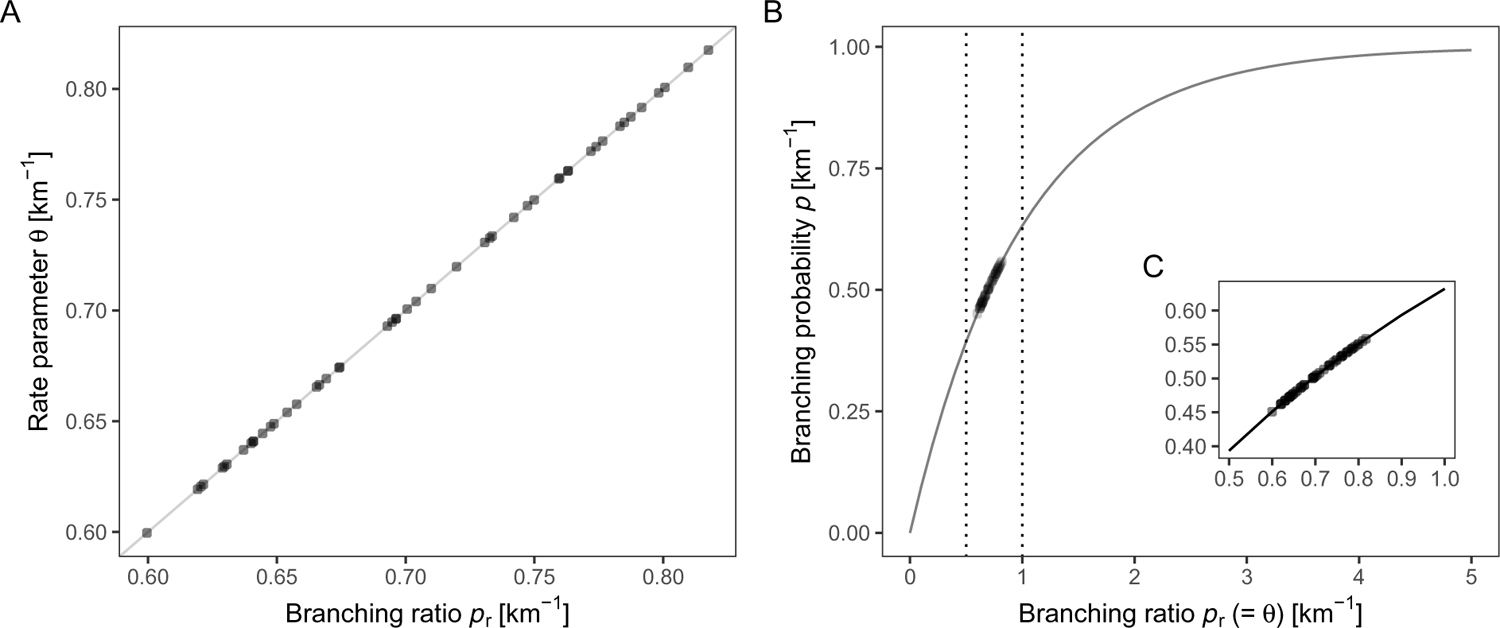
(A) Equality between rate parameter *θ* and branching ratio *p_r_*. Dots indicate estimated values at 50 rivers analyzed in Carraro and Altermatt^1^. The gray line denotes a 1:1 relationship. (B) Relationship between branching probability (*p* = 1 – exp(–*θ*)) and ratio 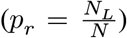. Dots are estimated values of *p* and *p_r_* at the 50 rivers. Note that *p* and *p_r_* were estimated independently; nevertheless, the data points fell exactly on the theoretical relationship between *p* and *θ* (the gray line), confirming the equality between *θ* and *p_r_*. Inlet zooms into the observed range of *p_r_*, which is denoted by the vertical broken lines.

Second, the term “scale invariance” was falsely used in their article. To explain this, let me consider an object *y* whose structural property is assessed under observation scale *x*. For example, Mandelbrot^10^ studied the length of a coastline as multiples of a ruler with a unit length *x* (the “observation scale”). In this practice, the observed object is expressed as a function of scale *x* (*y* = *f*(*x*)), and the function *f*(*x*) is said to be scale invariant if it suffices the following equation^11,12^:

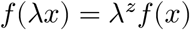

where *λ* is an arbitrary constant and *z* is the scaling exponent. The above equation is interpreted as the sign of “scale invariance” because the multiplicative extension/shrink of observation scale *x* by factor λ results in the same shape of the original object *y* but with a *different scale*^12^ (i.e., the structural property is retained). The power-law function is the most famous example that suffices this definition of scale invariance^11,13^:

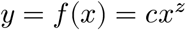

where *c* is the scaling constant. The scale invariance can be easily proved by multiplying the scaling factor λ:

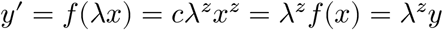

Carraro and Altermatt^1^ provided evidence that branching ratio *p_r_* follows a power law along the axes of observation scale *A*_*T*_ and pixel size *l* (i.e., length on a side) using OCNs (Equation 1 in the original article). Also, the authors visually showed that the relationship between *p_r_* and *A*_*T*_ follows a power-law using the data from 50 real river networks (Figure 3a in the original article). Nevertheless, the authors claimed that “*Here we show that an alleged property of such random networks (branching probability) is a scale-dependent quantity that does not reflect any recognized metric of rivers’ fractal character*…” (Abstract) This interpretation is the complete opposite as it has been defined in the literature of scale-invariance^11–14^, including the author’s previous publication^15^. Instead, by definition, the conclusion derived from their analysis must be “*branching ratio is a scale invariant feature that reflects the fractal nature of rivers.”*

Third, dimensions and units are improperly treated in their article. A dimension is the power of an axis along which a physical quantity is measured, and a unit is a way to assign a number to a particular dimension to make it relative. For a simple straight line, length is a dimension (dimension = 1), and a meter is a unit of length. Throughout the article, the authors used the number of pixels to measure the total river length *N*, the total catchment area *A*, and the threshold catchment area *A_T_*. There is no issue with using the number of pixels as a unit. However, a problem in their article is that they obscured the dimensions of pixels. For example, they made it very unclear that the river length *N* has a dimension of pixel “length,” while the total catchment area *A* and the threshold catchment area *A_T_* have a dimension of pixel “area.” In particular, the authors incorrectly defined *p_r_* as a dimensionless quantity (Methods) even though its unit is [pixel length^−1^].

The improper use of units caused more serious problems in the analysis of “real” river networks; the authors used different pixel sizes (length on a side) among watersheds (103 m to 1268 m; Supplementary Table 2 in the original article), meaning that the same number of pixels translates into very different lengths and areas. For example, in Figure 3 in the original article, the observation scale *A_T_* ranges 20 – 500 pixels. This pixel range translates into 0.2 – 5.3 km^2^ for the Toss river (with smallest pixel size) and 32.2 – 803.9 km^2^ for the Stikine river (with largest pixel size). Further, the branching ratio *p_r_* is also affected by this variation in pixel size as its unit is [pixel length^−1^]. Once the pixel length is converted to a unit of km, the branching ratio represents the number of links per 0.1 km in Toss, whereas it represents the number of links per 1.3 km in Stikine. Hence, the authors compared incomparable values, and their results reported in their article are not reliable.

To explore the consequences of the improper use of pixels, I re-analyzed their data with MERIT Hydro^9^; as noted above, this layer has constant pixel size, unlike the author’s analysis. I extracted river networks with 20 values of *A_T_* [km^2^] (*A_T_* = 1, …, 1000 with an equal interval at a log_10_ scale, but confined to *A_T_* < *A* for small rivers), at which I estimated branching ratio as 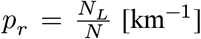. In the original article, the authors did not perform any statistical analysis even though *p_r_* (= *θ*) is a statistical parameter characterizing a link length distribution (see above); thus, its estimation accuracy is affected by the number of links, i.e., the sample size. To statistically substantiate the power-law of *p_r_* along the axis of *A_T_*, one must account for this heterogeneity of estimation accuracy. To this end, I fitted the following log-linear models with robust regression (*i* represents an individual data point of *p_r_* estimated in a given watershed at a given scale *A_T_*).

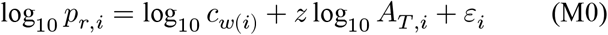

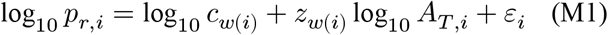

where *ε*_*i*_ is the error term that is properly weighted by Huber’s function. Robust regression analysis is appropriate because it is robust to outliers caused by the small sample size (typical for large *A_T_* values). The first model (M0) assumes the “universal” scaling with the single exponent *z* across watersheds; i.e., the branching ratio at all the 50 watersheds follows the same power law with the watershed-specific constant log_10_ *c*_*w*(*i*)_ (*w*(*i*) is watershed *w* for a data point *i*). In contrast, the second model (M1) assumes the “localized” scaling with the watershed-specific exponent *z*_*w*(*i*)_. I estimated the evidence ratio of the two models using the approximated Bayes Factor (BF)^16^, which is defined as 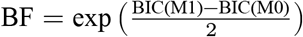. In this definition, a value of BF > 1 gives the support for M0 over M1; for example, if BF = 2, the model M0 is twice as likely as the model M1.

The analysis provided decisive support for M0 with BF ≈ 3.00×10^20^. Under M0, as evident from its model formula, the rank of the expected branching ratio E(log_10_ *p_r_*) never changed across scales, maintaining the order of the watershed-specific constant log_10_ *c*_*w*(*i*)_ (Figure 2; see regression lines). This result is inconsistent with the author’s statement “*by extracting different river networks at various scales (i.e., various A*_*T*_ *values) and assessing the rivers’ rank in terms of p*_*r*_, *one observes that rivers that look more “branching” (i.e., have higher p*_*r*_*) than others for a given A*_*T*_ *value can become less “branching” for a different A*_*T*_ *value (Fig. 3)*.” I also must note that I did not find any significant correlation between watershed area *A* [km^2^] and branching ratio *p_r_* [km^−1^] when extracted with a constant value of *A_T_* = 1 km^2^ across watersheds (Spearman’s rank correlation = –0.19, p-value = 0.19), as opposed to the statement in the original article “…*if different river networks spanning different catchment areas (say, in km*^2^*) are compared, all of them extracted from the same DEM (same l and same A*_*T*_ *in km*^2^*), then the larger river network will appear more branching (i.e., have larger p*_*r*_)^1^.” The lack of correlation between branching probability *p* and watershed area *A* has also been reported in two previous studies^6,7^, one of which used 184 watersheds for the analysis^7^. This result is not surprising at all because *p_r_* is the number of links “divided” by the total river length by definition. My re-analysis revealed that the author’s statements merely reflect the lack of appropriate quantitative analysis and/or a statistical artifact of inconsistent units across watersheds.

**Figure 2:**
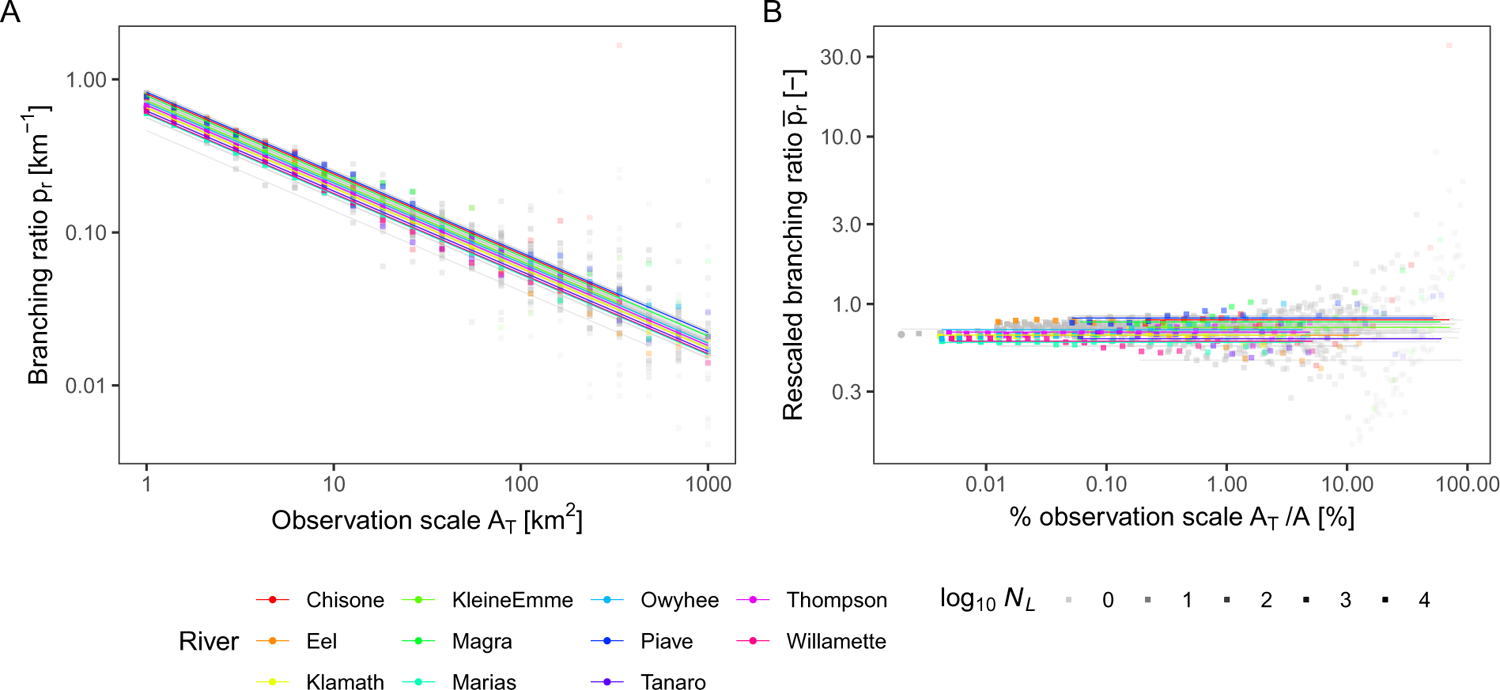
Log­log plot substantiates the power­law scaling of dimensional branching ratio (*p_r_*; Panel A) along the axis of observation scale *A_T_*. Once non­dimensionalized, the properly rescaled branching ratio (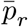; Panel B) will converge to values unique to each river as *A_T_* → 0. Colors indicate rivers highlighted in the original article^1^. Individual data points are shown in dots, whose transparency is proportional to the number of links *N*_*L*_ (i.e., sample size). Lines are predicted values from the model M0 (i.e., the expected value of dimensional or rescaled branching ratio).

It is worthwhile to note that if scaling exponent varies by watershed (i.e., *z*_*w*_; albeit not the case for my analysis), the dimensional *p_r_* is incomparable across watersheds because its dimension (= 2*z*_*w*_) is different (just like length and area are not comparable). To understand this argument, one must recognize that, by writing 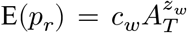, we equate the dimension in both sides of the equation. From this equation, the dimension of *p_r_* is “estimated” to be 2*z*_*w*_ since the scaling exponent *z*_*w*_ applies to the unit of *A_T_* as 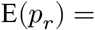 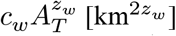. Therefore, to compare branching ratio across watersheds, we must non-dimensionalize *p_r_* as 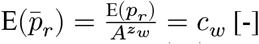 (see Figure 2B for this exercise). Evidently, the expected value of dimensionless branching ratio 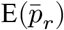 remains constant across scales (*c*_*w*_) with no possibility of changes in the rank (Figure 2B). This technique of dimensional analysis has been widely used when comparing or non-dimensionalizing the self-similar structure of scale-invariant objects (see Figure 1 in Rinaldo et al.^14^ and equation (2.2) in Rodriguez-Iturbe and Rinaldo^11^ for examples). In fact, past empirical studies used *A_T_* = 1 km^2^ such that the estimated dimensional *p_r_* approximates the rescaled 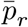 (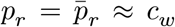 for *A_T_* = 1 in a given unit, regardless of the value of *z*)^6,7^. Note that 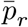 fluctuates unpredictably at coarser observation scales (larger *A_T_* relative to *A*; Figure 2B). This is simply because of the small sample size (i.e., the number of links) at coarser resolutions, which inflates the statistical uncertainty of parameter estimation for *p_r_*. However, a solution is simple: use a sufficiently small observation scale (e.g., *A_T_* = 1 km^2^) to yield a large sample size *N*_*L*_. The 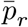 will converge to the single value that represents the self-similar structure unique to each river network (Figure 2B). Therefore, there is no mathematical supporting ground for the author’s statement “…*branching probability is a non-descriptive property of a river network, which by no means describes its inherent branching character*…”

Lastly, the issue of units is pertinent to their metapopulation simulations, as they used pixel size *l* as a unit of “population scale” and “dispersal distance.” This makes their simulation results hardly interpretable. For example, they assumed a population scale of 103 m in Toss, while assuming 1268 m in Stikine. A reasonable interpretation of this simulation setup is that the authors simulated metapopulation dynamics of “different” species among watersheds. This is not trivial as the population scale defines the number of local populations within a metapopulation – one of the most influential parameters dictating metapopulation CV (see Equations (2) and (3) in the original article). If one assumes the same population scale across watersheds (i.e., the same species), then Stikine should be ~12 times greater in the number of local populations *N*_*p*_ compared to Toss. Nevertheless, the authors used a nearly-constant number of *N*_*p*_ for all the 50 real rivers (e.g., E(*N*_*p*_) ≈ 1088 for *A_T_* = 500 [pixel area]) by changing the population scale. Similarly, the average dispersal distance *α* was measured in the unit of pixel length (*α* = 100 [pixel length] in Figure 6 in the original article), meaning that it varies from 10.3 km to 126.8 km among watersheds when converted to a unit of km. It is difficult to envision that such a huge variation in dispersal capability exists within the same species. Again, the parameter *α* has a decisive influence on metapopulation CV and capacity in the author’s model (see Methods in the original article). Critically, Figure 6 and Supplementary Figures 4 – 9 of the original article “aggregated” these metapopulations of different species in real rivers and compared the overall summary statistics (e.g., median) with those obtained from BBTs, RBNs, and OCNs. Therefore, the authors compared incomparable values. A better alternative should have been to use the same pixel size (= population scale) among watersheds to assume metapopulations of the “same” species, then pairing metapopulation metrics by watershed. In doing so, the authors could examine how predicted values from different virtual networks correlate with values in real rivers with a reference to a 1:1 line. Although I do not know how this improvement will alter one of the main conclusions “*OCNs most accurately predicted the metapopulation stability and capacity*,” it is clear that their simulation results are not ecologically interpretable.

## Data availability

Codes and data are available at https://github.com/aterui/public-proj_fractal-river.

